# Periaqueductal gray passes over disappointment and signals continuity of remaining reward expectancy

**DOI:** 10.1101/2024.12.17.628983

**Authors:** Hyunchan Lee, Okihide Hikosaka

## Abstract

Disappointment is a vital factor in the learning and adjustment of strategies in reward-seeking behaviors. It helps them conserve energy in environments where rewards are scarce, while also increasing their chances of maximizing rewards by prompting them to escape to environments where richer rewards are anticipated (e.g., migration). However, another key factor in obtaining the reward is the ability to monitor the remaining possibilities of obtaining the outcome and to tolerate the disappointment in order to continue with subsequent actions. The periaqueductal gray (PAG) has been reported as one of the key brain regions in regulating negative emotions and escape behaviors in animals. The present study suggests that the PAG could also play a critical role in inhibiting escape behaviors and facilitating ongoing motivated behaviors to overcome disappointing events. We found that PAG activity is tonically suppressed by reward expectancy as animals engage in a task to acquire a reward outcome. This tonic suppression of PAG activity was sustained during a series of sequential task procedures as long as the expectancy of reward outcomes persisted. Notably, the tonic suppression of PAG activity showed a significant correlation with the persistence of animals’ reward-seeking behavior while overcoming intermittent disappointing events. This finding highlights that the balance between distinct tonic signaling in the PAG, which signals remaining reward expectancy, and phasic signaling in the LHb, which signals disappointment, could play a crucial role in determining whether animals continue or discontinue reward-seeking behaviors when they encounter an unexpected negative event. This mechanism would be essential for animals to efficiently navigate complex environments with various reward volatilities and ultimately contributes to maximizing their reward acquisition.

## Introduction

During reward-seeking behaviors, humans and animals occasionally encounter disappointing events that diminish their engagement and even prompt them to leave the environment. Disappointment could be a crucial factor, for example, in environments where resources become scarce, helping animals adjust their behavioral patterns to minimize unnecessary energy expenditure (e.g., by hibernating) or migrate to environments with richer resources.^1–6^ However, in real life, we often need to pass through a series of unexpected events, some of which may be disappointing or irrelevant to immediate gratification, to achieve our desired outcomes ultimately.^7^ Thus, we hypothesized that there would be another important neuronal mechanism that signals the remaining reward expectancy, helping us overcome disappointing events and continue motivated behaviors until we reach our desired goals.^8–10^ Then, how does the animal brain signal disappointment and the remaining reward expectancy when encountering a disappointing event?

To solve this question, we examined the lateral habenula (LHb) and periaqueductal gray (PAG), key brain regions involved in the regulation of negative mood in animals.^11–13^ The LHb is a well-known primary input to dopamine neurons, especially signaling disappointment^14^ and aversiveness^15^ from stimuli. Thus, dysfunction of the LHb has been extensively studied as a promising therapeutic target for treating major depressive disorder.^16–19^ In the present study, we raised a question regarding a feature of the LHb response, which primarily involves phasic firing lasting 100-500 ms.^20^ This rapid signaling is crucial for animals to quickly learn and identify significant objects in a complex environment, where various valued objects are presented sequentially, and to promptly guide their actions step-by-step accordingly.^21–24^ However, if an animal’s reward-seeking behavior, which involves a series of sequential actions, is solely regulated by the phasic signaling of the LHb, the animal would easily abandon or leave the ongoing motivated behavior every time it encounters a disappointing event.

Therefore, we investigated another key brain region involved in the regulation of negative moods, the PAG.^25–27^ Notably, PAG activity has primarily been reported to play a pivotal role in controlling coping strategies in response to aversive stimuli, such as defensive and escape behaviors.^28^ As a result, the PAG has been suggested to be a critical brain area that can provide therapeutic interventions not only for mental disorders^29–31^ but also for movement disorders related to the control of involuntary behaviors.^32–37^ This has been further supported by previous studies showing that PAG manipulation affects pain, analgesia,^38^ uncontrollable freezing behaviors,^39^ and involuntary vocalizations.^40^ Moreover, the PAG has crucial efferents that project to the motor control system, such as the raphe interpositus, which regulates gaze fixation in primates.^41^ Hence, we could expect that the PAG would not only trigger escape behaviors but also play a critical role in controlling escape and sustaining reward-seeking behaviors when an animal encounters a disappointing event.

In summary, we investigated how the LHb and PAG neurons signal disappointment and the remaining reward expectancy to understand how an animal decides whether to overcome or leave a disappointing event. As a result, we found a significant correlation between PAG activity and the continuity of reward-seeking behavior in animals, which could help them overcome disappointing events and ultimately obtain their desired rewards.

## Results

To record the neuronal and behavioral responses of rhesus macaque monkeys while passing through a series of disappointing events before ultimately obtaining a reward outcome, we devised a scene-based foraging/Pavlovian task. In this task, two different foraging and Pavlovian tasks were conducted on a shared background scene image (Fig. 1A). As a result, the monkeys were able to predict different reward outcomes for each group of scene images based on the average reward outcomes experienced in the foraging and Pavlovian tasks. Therefore, the monkeys experienced multiple changes in reward predictions throughout the task, depending on the appearance of each group of scene images and tasks.

**Figure 1.**
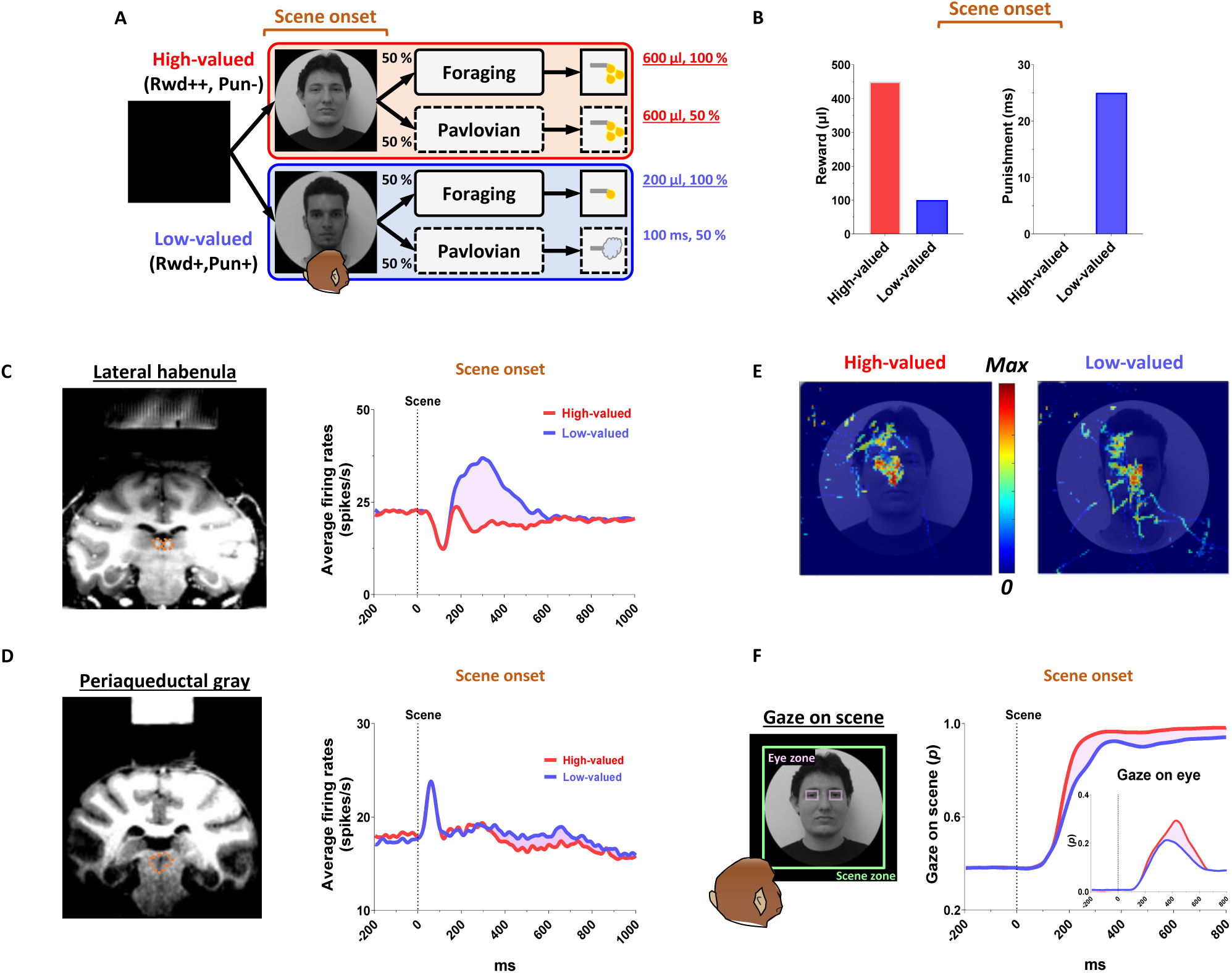
Neuronal responses and anticipatory gazes in scene viewing. (A) In the scene-based foraging/Pavlovian task, each trial began with the free viewing of a scene image. Each group of scene images provided respective reward outcomes from the foraging and Pavlovian tasks. Consequently, the monkeys freely viewed each group of scene images based on the average rewards they had experienced during both tasks. (B) Midlines indicate the average amount of juice reward and airpuff punishment provided to each group of scenes from the foraging and Pavlovian tasks. Bars indicate the maximum and minimum amounts of the outcomes. (C, D) Left, recording sites are marked with orange dotted lines, indicating the locations of the LHb and PAG. Right, the average firing rates of LHb and PAG in response to scene onset. The purple shaded areas represent the differences in neuronal responses between high-valued and low-valued scenes. (E) An example heatmap illustrating monkeys’ gaze on face scene images. (F) Outside, the probabilities of gaze on face scene images during the free-viewing period (Scene zone, 40° × 40°). Inside, the probabilities of gaze on the eye regions of face scene images during the free-viewing period (Eye zone, 4° × 3°). Please see Supplementary Fig. 1 for additional information and *P* values.

### Reward expectation facilitates visual attention to contexts

We firstly found that reward experiences facilitate the visual attention of primates. Each trial of the task started with the appearance of a scene image (Fig. 1A, scene onset), which allowed the monkey’s free-viewing for 1 s and then remained as a background scene until the end of the trial, during which either the Pavlovian or foraging task was performed. The monkeys experienced large amount of juice rewards in both foraging and Pavlovian tasks in the high-valued scenes (Fig. 2A and C), while they received small amount of juice rewards in the foraging task and airpuff punishments in the Pavlovian task in the low-valued scenes (Fig. 2B and D). As a result, at the start of a trial, monkeys could predict greater reward outcomes from the appearance of the high-valued scenes compared to low-valued scenes (Fig. 1B).

**Figure 2.**
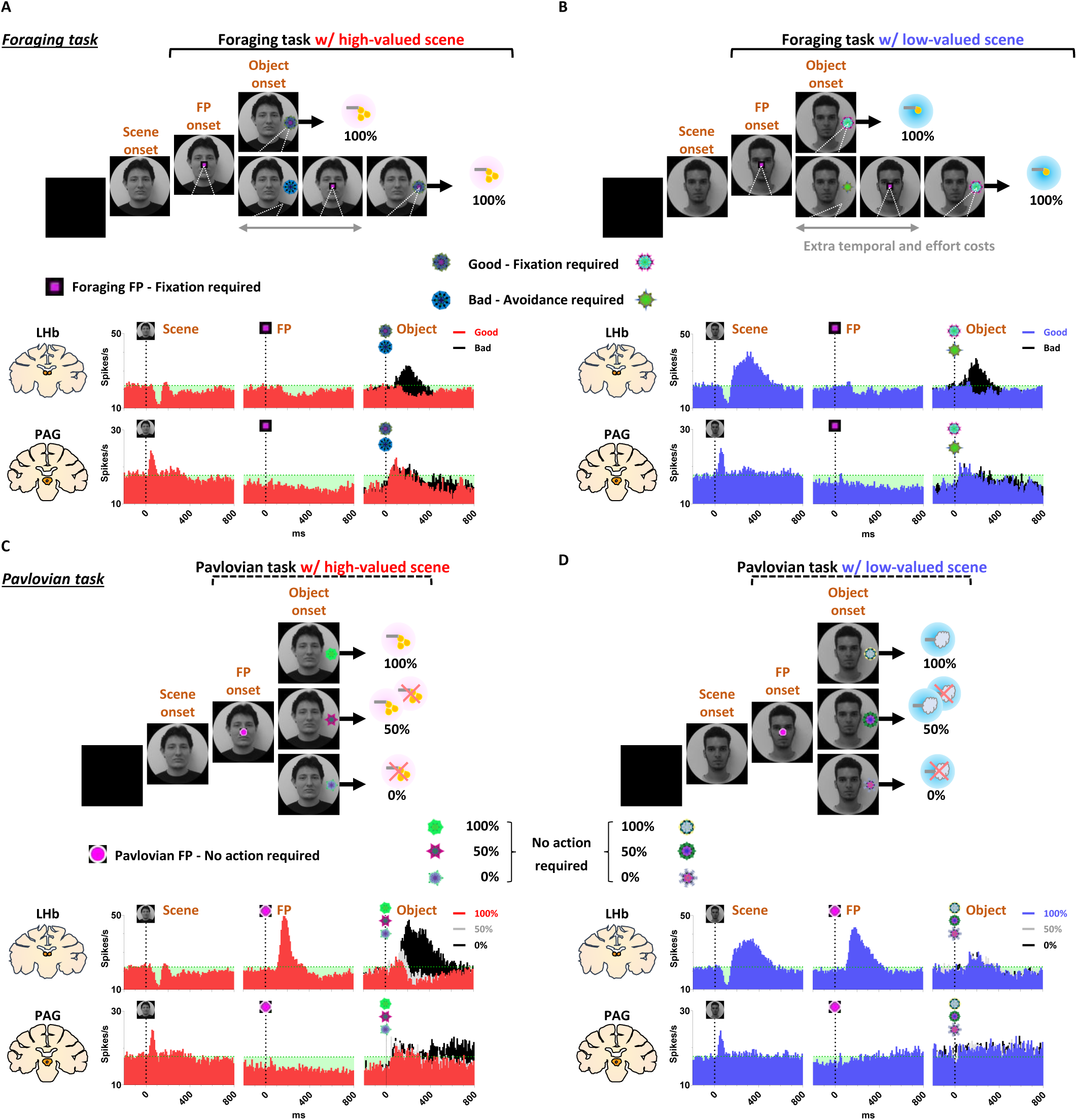
Neuronal and behavioral responses in foraging and Pavlovian tasks. (A) The histograms (10 ms bins) showing neuronal activity in the LHb (top) and PAG (bottom) during the foraging task with high-valued scenes. The histograms are aligned to the onset of each stimulus during the task procedures (scene onset, FP onset, and object onset). The green dotted line indicates baseline neuronal activity measured for 200 ms before scene onset. The green shaded areas represent changes in neuronal activity compared to baseline levels during the task procedures. (B) The histograms show the same data as in A, but for low-valued scenes. (C-D) The histograms show the same data as in A and B, respectively, for the Pavlovian task.

As the monkeys experienced larger rewards, their free viewing became more focused on high-valued scenes (Fig. 1F, outside). For the background scene images, we used face and landscape images, as they are representative examples of social and spatial contexts that animals encounter during reward-seeking behaviors in real life.^42^ When we used face images as the background scene images, they showed more pronounced gazes toward the eye regions in the high-valued face scene images than in the low-valued scene images, accompanied by stronger scene viewing of the high-valued face images (Fig. 1E and F, inside).

Through an additional experiment where the influence of reward and punishment on scene gaze was examined respectively, we confirmed that the effect of value on monkeys’ gaze for scene images was particularly regulated by the predicted values of reward rather than airpuff punishment in this task design (Supplementary Fig. S1). The monkeys’ gaze toward the scene images was more focused on those with expected high-value rewards, even if they included an airpuff punishment, and it decreased for images with expected low-value rewards, even if there was no airpuff punishment.

### Continuity of reward expectancy facilitates subsequent actions even after disappointment

We then found that the reward expectancy from the scene images continuously facilitated subsequent actions of animals in the next step. As described above, each trial began with the appearance of a scene image and the monkeys’ free viewing for 1 s (Fig. 1A). In the meantime, although the monkeys’ gaze was more focused on the high-valued scenes than on the low-valued scenes, we found that their gaze toward the scene images commonly increased during the 50-150 ms period for both scenes, reaching over 90% and being sustained until the next stimuli appeared (Fig. 1F, outside). At the onset of the low-valued scenes, the monkeys sometimes closed their eyes or shifted their gaze away from the scene image during the free viewing period. However, they soon returned their gaze to the scene and prepared to perform the subsequent task. This indicates that the monkeys had a strong motivational engagement for the subsequent task procedures in both scene groups. Notably, these highly motivated states in both scenes consistently facilitated subsequent behaviors.

After a free-viewing of the scene for 1 s, a fixation point (FP) appeared at the center of the screen, and either the foraging task or the Pavlovian task was initiated (Fig. 2). The foraging task and Pavlovian task were distinguished by different shapes of the FP (Fig. 2A-B, foraging task, square; Fig. 2C-D, Pavlovian task, circle). Once the FP appeared at the center of the scenes, the monkeys quickly fixated their gaze on the FP within 20 ms, whether it indicated the foraging task or the Pavlovian task (Fig. 3D and H). In both foraging and Pavlovian tasks, the start time at which monkeys’ gaze reached the FP was significantly quicker in high-valued scenes than in low-valued scenes. In addition to the speed of the fixation start time, the monkeys exhibited higher fixation rates in high-valued scenes compared to low-valued scenes (Fig. 3C and G). However, even in low-valued scenes, monkeys initiated fixation on the FP within 20 ms (Fig. 3D and H, blue), and the fixation rates were over 95% (Fig. 3C and G, blue). This implies that the monkeys were in a strong motivational state in both high-valued and low-valued scenes. Indeed, even before the FP onset, about 90% of the monkeys’ gaze had already stayed on the location where the FP would appear (Fig. 3 A-B and E-F).

**Figure 3.**
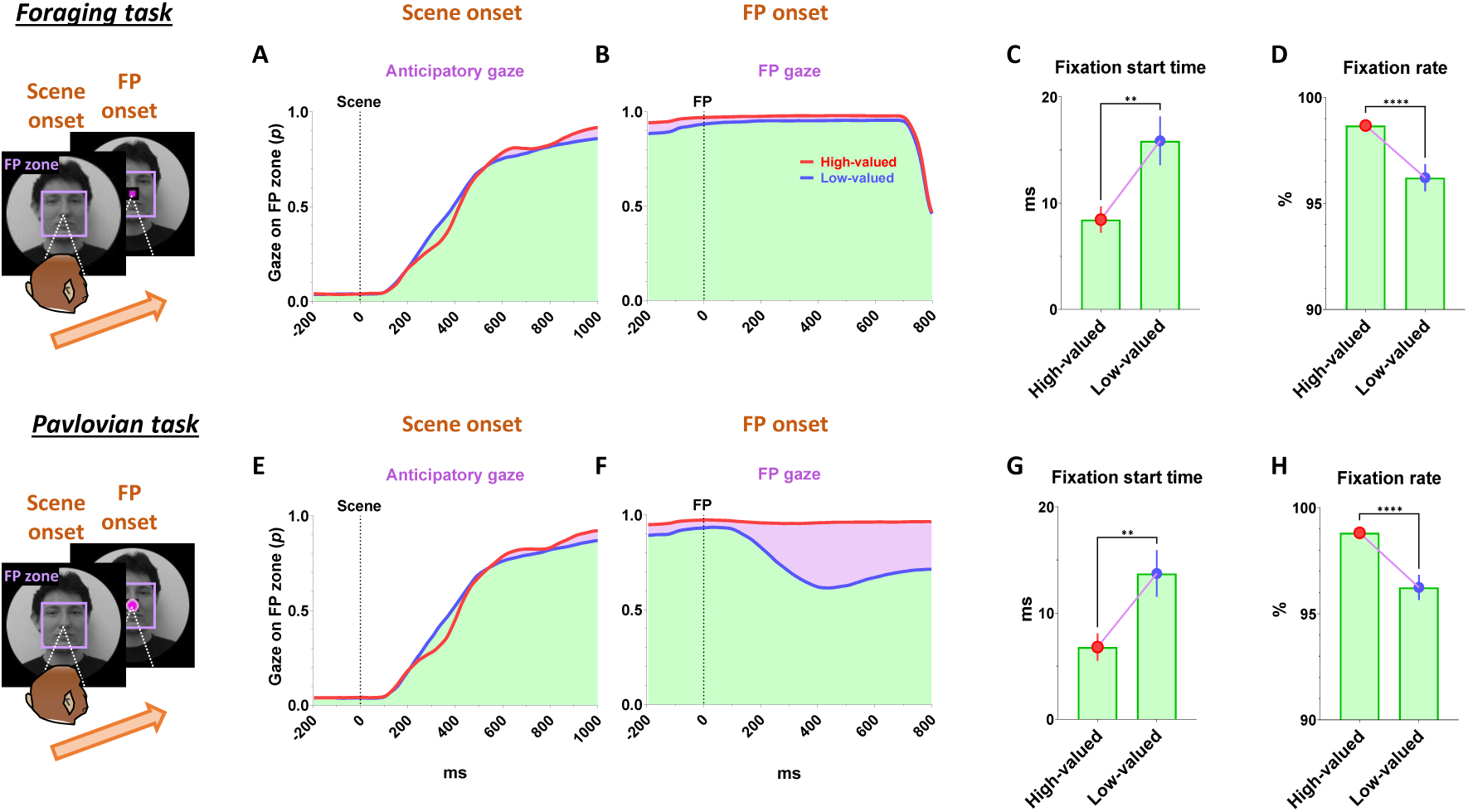
Reward expectancy facilitates subsequent actions. (A) The probabilities of anticipatory gaze before the FP onset, where the gaze was located at the position where the FP would appear (FP zone, 10°× 10°). The purple shaded areas represent differences in responses between high-valued and low-valued scenes or between good objects and bad objects. The green shaded areas indicate the monkeys’ gaze on the location where the FP appeared, starting from scene onset and maintained throughout the subsequent task procedures. The purple shaded areas highlight the differences in gaze responses between high-valued and low-valued scenes. (B) The probabilities of gaze on the FP after its appearance. (C) The fixation start time, representing the gaze-reaching time for monkeys to initiate fixation on the FP (Wilcoxon matched-pairs signed rank test; *\*\*P* < 0.01, *n* = 72). (D) The fixation rate, indicating the percentage of time monkeys maintained their gaze on the FP to engage in the next task procedure (Wilcoxon matched-pairs signed rank test; *\*\*\*P* < 0.0001, *n* = 72). (E-F) Same as A-B, respectively, for the Pavlovian task. (G-H) Same as C-D, respectively, for the Pavlovian task.

This result implies that as long as reward expectancy is sustained, animals can maintain a strong motivational state even after disappointment. How, then, do the neural mechanisms in the brain operate to sustain such a strong motivational state in animals, even after the disappointment?

### PAG neurons signal reward expectancy using tonic activity

We found that the neuronal activity of the PAG was tonically modulated by the expectancy of reward outcomes. Along with the monkeys’ gaze behaviors, we recorded neuronal activities in the LHb and PAG (Fig. 2). When a scene image appeared at the beginning of a task trial, neuronal activity in both brain regions was inhibited by high-valued scenes and excited by low-valued scenes (Fig. 1C and D). However, their responses exhibited distinct characteristics, with the PAG primarily showing tonic firing, while the LHb responded with phasic firing. We notably found that the reward expectancy induced by the scene onset had a significant and continuous effect not only on the continuity of the monkeys’ subsequent scene-viewing behaviors (Fig. 1F, outside) but also on the tonic activity of PAG neurons, which persisted from scene onset to FP onset (Fig. 1D).

The phasic LHb response was advantageous for rapidly encoding value information from the scene images, with a short latency of around 150 ms (Fig. 1C). The neuronal activity of LHb was suppressed by high-valued scenes and excited by low-valued scenes. The difference between these two scenes persisted for about 450 ms, after which their firing rates quickly recovered to baseline levels of neuronal activity, similar to those before the onset of the scene. This enabled the LHb to cease its response to the scene images before the next stimuli appeared and to prepare for subsequent responses in the following steps (Fig. 4J and 5J, before 0 ms).

**Figure 4.**
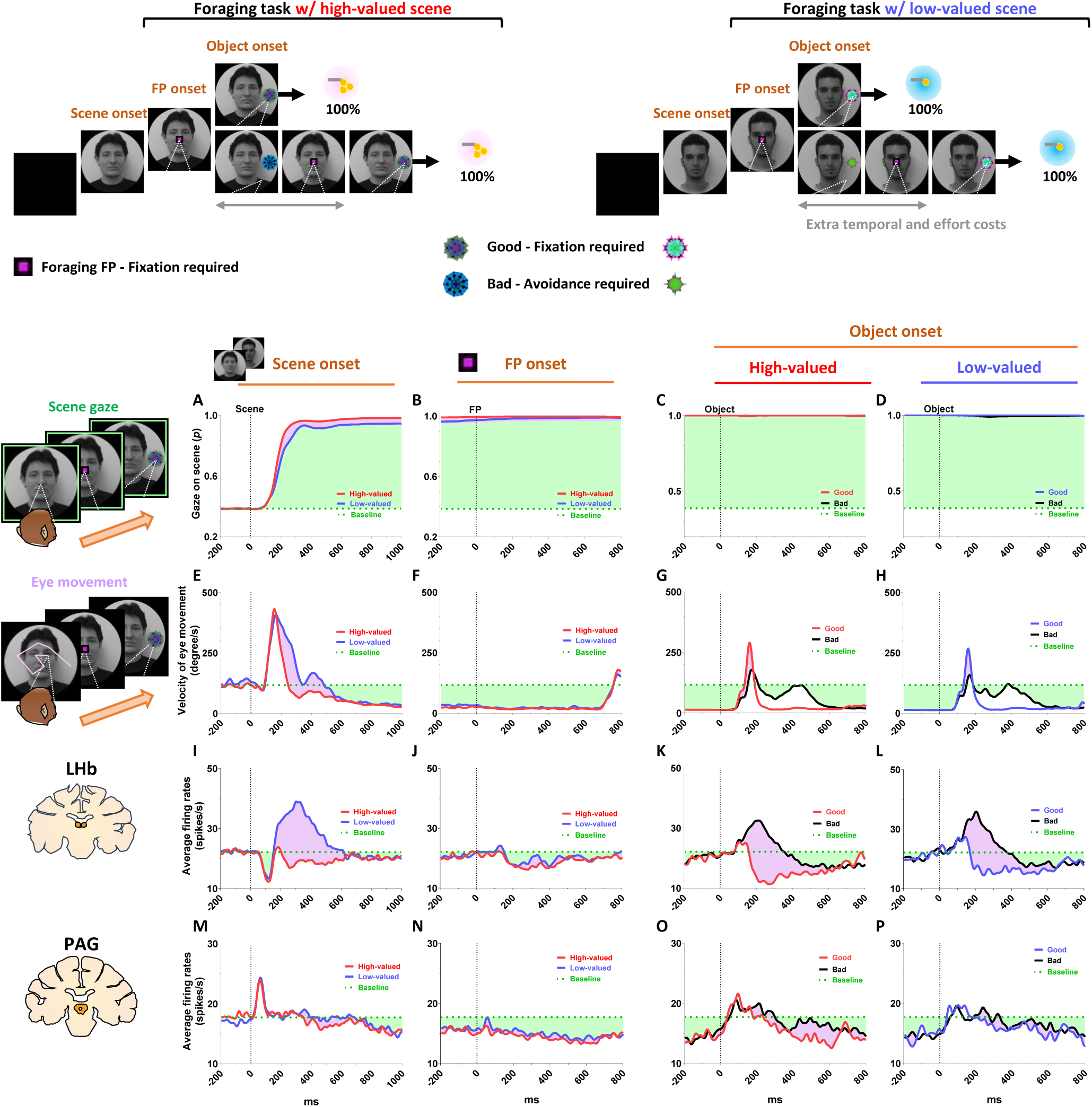
Tonic inhibition of PAG signals continuity of reward expectancy in foraging task. (A-D) The probabilities of gaze fixation on scene images during the foraging task. The purple shaded areas represent differences in responses between high-valued and low-valued scenes, or between 100% objects and 0% objects. The green dotted line indicates baseline levels measured for 200 ms before scene onset, while the green shaded areas represent changes in gaze responses relative to baseline during the task procedures. (E-H) Same as A-D, but for the velocity of eye movement. (I-L) The LHb activity shown in figures 2A-B aligned with the gaze response presented in A-H. (M-P) The PAG activity corresponds to the data presented in I-L.

**Figure 5.**
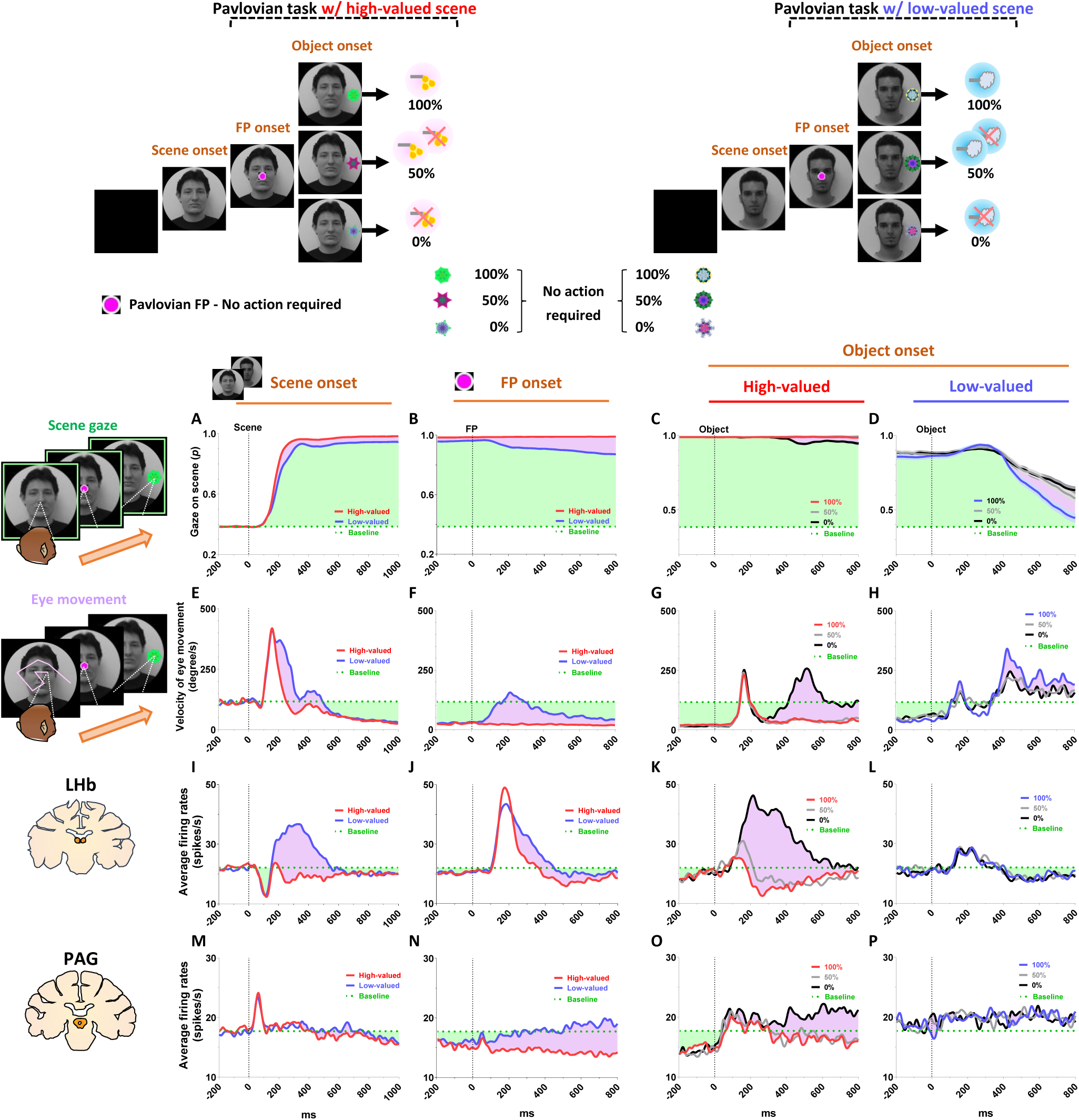
Tonic excitation of PAG signals complete extinction of reward expectancy in Pavlovian task. (A-D) The probabilities of gaze fixation on scene images during the Pavlovian task. The purple shaded areas represent differences in gaze responses between high-valued and low-valued scenes. The green dotted line indicates baseline levels measured for 200 ms before scene onset, while the green shaded areas represent changes in gaze responses relative to baseline during the task procedures. (E-H) Same as A-D, but for the velocity of eye movement. (I-L) The LHb activity shown in figures 2C-D aligned with the gaze response presented in A-H. (M-P) The PAG activity corresponds to the data presented in I-L.

In contrast, the tonic activities of PAG slowly encoded the value information provided by the scene onset, starting around 300 ms (Fig. 1D). The neuronal activity of the PAG was also inhibited in response to high-valued scenes and slightly excited in response to low-valued scenes. However, after the value responses, the PAG activity did not quickly recover to baseline levels as the LHb responses did. Meanwhile, the PAG responses exhibited a more pronounced tendency to steadily sustain the tonic suppression in both high-and low-valued scenes until the next visual information was presented (Fig. 4N and 5N, before 0 ms). This result implies that PAG activity could tonically regulate the persistence of the task-engaged state in animals from a reward-informative stimulus to a subsequent reward-informative stimulus.

### Tonic PAG activity passes over disappointment and signals the continuity of reward expectancy until the goal is ultimately obtained

We found that the PAG response is specifically tuned to signal the continuity of reward expectancy, even when disappointment occurs during the sequential procedures of reward-seeking behaviors. As described above, the PAG neurons exhibited tonic suppression despite the disappointment induced by the low-valued scene (Fig. 1D). Especially through the foraging task, we observed additional evidence showing that PAG activity was able to maintain tonic suppression despite further disappointments occurring, until the reward was ultimately obtained (Fig. 2A-B; Fig. 4M-P).

The foraging task was designed to observe behavioral and neural responses when temporary disappointment occurs during sequential task procedures, but the possibility of obtaining a reward outcome is 100% guaranteed. The foraging task began with the appearance of the square-shaped FP on the scene images. Once the monkeys completed gaze fixation on the FP, the FP disappeared, and good or bad objects appeared randomly on the left or right side (Fig. 2A-B, Object onset). The appearance of the good object allowed the monkeys to receive a reward immediately upon completing gaze fixation on it (Supplementary Movie S1). In contrast, the bad object required avoiding gaze toward it (Supplementary Movie S2). The monkeys could avoid the bad object by either not making a saccade toward it for 1 s or making a saccade but breaking the fixation within 500 ms. After the monkeys successfully avoided gazing at the bad object, the FP reappeared. Subsequently, the monkeys were finally able to find the good object and received a reward by fixating on it. The monkeys performed these instrumental behaviors with a success rate of over 95%. Even if they failed, the same trial was repeated until they succeeded, ultimately receiving a reward outcome with a 100% probability.

On the one hand, we found that the phasic response of the LHb was proficient in encoding the value difference between good and bad objects (Fig. 4K-L). The appearance of the bad object was another disappointing event for the monkeys, as it incurred additional temporal and effort costs compared to the good object. As a result, LHb activity was inhibited by the presence of good objects and excited by the presence of bad objects.

On the other hand, we found that the tonic activity of the PAG maintained its tonic suppression state throughout the sequential procedures of the foraging task, even when good or bad objects appeared, ultimately leading to a 100% probability of obtaining a reward outcome (Fig. 4O-P). We observed that the tonic suppression of the PAG, which began with the initial scene onset (Fig. 4M), was sustained until the FP onset (Fig. 4N) and continued even after either the good or bad objects appeared (Fig. 4O-P). Although PAG activities were slightly excited by the appearance of either good or bad objects, their activities quickly declined and returned to a level lower than baseline (Fig. 4O-P, before 0 ms). As a result, the firing rates of the PAG neurons remained lower than baseline levels when FP reappeared following the avoidance of the bad object.

### Distinct LHb and PAG responses signal disappointment and continuity of remaining reward expectancy

On the contrary, we also confirmed that PAG activity becomes tonically excited when the reward expectancy is completely extinguished during an ongoing motivated behavior, and remains in a tonically excited state until the end of the trial. This was evident in the Pavlovian task, which dramatically altered the remaining reward expectancy, sustaining it in high-valued scenes and completely extinguishing it in low-valued scenes.

In the high-valued scenes, the Pavlovian task resulted in different probabilities of obtaining rewards (100%, 50%, 0%) depending on the object presented (Fig. 2C; Supplementary Movie 3). Ultimately, the appearance of the FP-indicating the Pavlovian task in the high-valued scenes resulted in an average probability of obtaining rewards of 50% (the average of 100%, 50%, and 0%). This implies that it caused disappointment compared to the 100% probability of obtaining rewards in the foraging task (Fig. 2A), but there is still a possibility of obtaining rewards.

On the other hand, in the low-valued scene, when the Pavlovian task began with the appearance of the FP, the probability of obtaining rewards was completely extinguished to 0%, leaving only the prediction of punishment (Fig. 2D; Supplementary Movie 4). This not only caused disappointment compared to the 100% probability of obtaining rewards in the foraging task, but also implies that the possibility of obtaining rewards within the trial was completely extinguished (Fig. 2B and D).

As a result of the common disappointment in both high- and low-valued scenes, LHb activities were excited by the FP-indicating the Pavlovian task (Fig. 5J). However, PAG exhibited different patterns of excitation and inhibition in response to the FP, depending on whether the Pavlovian task was initiated in high-valued or low-valued scenes (Fig. 5N).

In the low-valued scenes, the PAG became excited when the expectancy of reward outcomes was completely extinguished by the appearance of the FP-indicating the Pavlovian task (Fig. 5N). Then, this tonically excited state of PAG activity was sustained until the end of the trial, remaining slightly above baseline levels when an object appeared after the FP (Fig. 5P, before 0 ms). The PAG activity was not responsive to variations in punishment probability (100%, 50%, or 0% airpuff), differentiated by the appearance of objects (Fig. 5P), but had no significant impact on the previously extinguished remaining reward expectancy.

In contrast, in the high-valued scenes, the Pavlovian task induced disappointment but still allowed the monkeys to maintain the expectancy of obtaining rewards, even though the probability was reduced. As a result, when the FP-indicating the Pavlovian task appeared in the high-valued scenes, the PAG remained in a suppressed state until an object appeared (Fig. 5O, before 0 ms). Subsequently, the neuronal activities of the PAG were significantly differentiated by the 100%, 50%, and 0% reward objects. When the objects associated with 100% or 50% reward probabilities appeared, the PAG activity remained in a tonically suppressed state, but it became excited in response to the 0% reward object, which also signifies the complete extinction of reward expectancy within that trial (Fig. 5O).

Consequently, we suggest that the distinct phasic LHb and tonic PAG responses could play separate roles in signaling disappointment and remaining reward expectancy. How, then, do changes in the tonic activity of the PAG contribute to behavioral responses when animals encounter an event that switches the remaining reward expectancy on or off?

### Tonic PAG activity facilitates subsequent actions based on inhibitory motor control

Finally, we propose that the tonic activity of PAG neurons could signal the continuity of remaining reward expectancy and facilitate subsequent actions based on inhibitory motor control. Our findings have so far revealed that PAG neurons exhibited tonic inhibition during the foraging task, which guaranteed a 100% probability of obtaining a reward (Fig. 4M-P). This tonic inhibition of the PAG corresponded to the continuity of reward expectancy, facilitating monkeys’ persistent visual attention throughout the task trials (Fig. 4A-D). Additionally, during this period, we observed that their overall eye movements were suppressed (Fig. 4E-H) while the PAG exhibited tonic inhibition. This inhibitory control of eye movements could reflect an increase in gaze fixation on the scene images, indicating the persistence of visual attention. Along with this, our further evidence suggests that the inhibitory control of eye movements could play additional crucial roles in facilitating the execution of subsequent actions.

This was evident in the Pavlovian task, which did not require any instrumental actions and allowed us to observe the natural behaviors of the monkeys. When the FP-indicating Pavlovian task appeared in low-valued scenes, the monkeys’ remaining reward expectancy was completely extinguished. As a result, PAG activity was tonically excited (Fig. 5N and P), and the monkeys exhibited an increase in eye movements (Fig. 5F and H), which in turn resulted in a decrease in scene gaze compared to high-valued scenes (Fig. 5B and D). Consequently, PAG activity (Fig. 5O-P, before 0 ms) and eye movements (Fig. 5G-H, before 0 ms) at the time of object onset were greater in low-valued scenes than in high-valued scenes (Fig. 6A). Ultimately, we found that PAG activity observed at the time of object onset was significantly correlated with the distance of the monkeys’ eye movements before the object onset (Fig. 6B).

**Figure 6.**
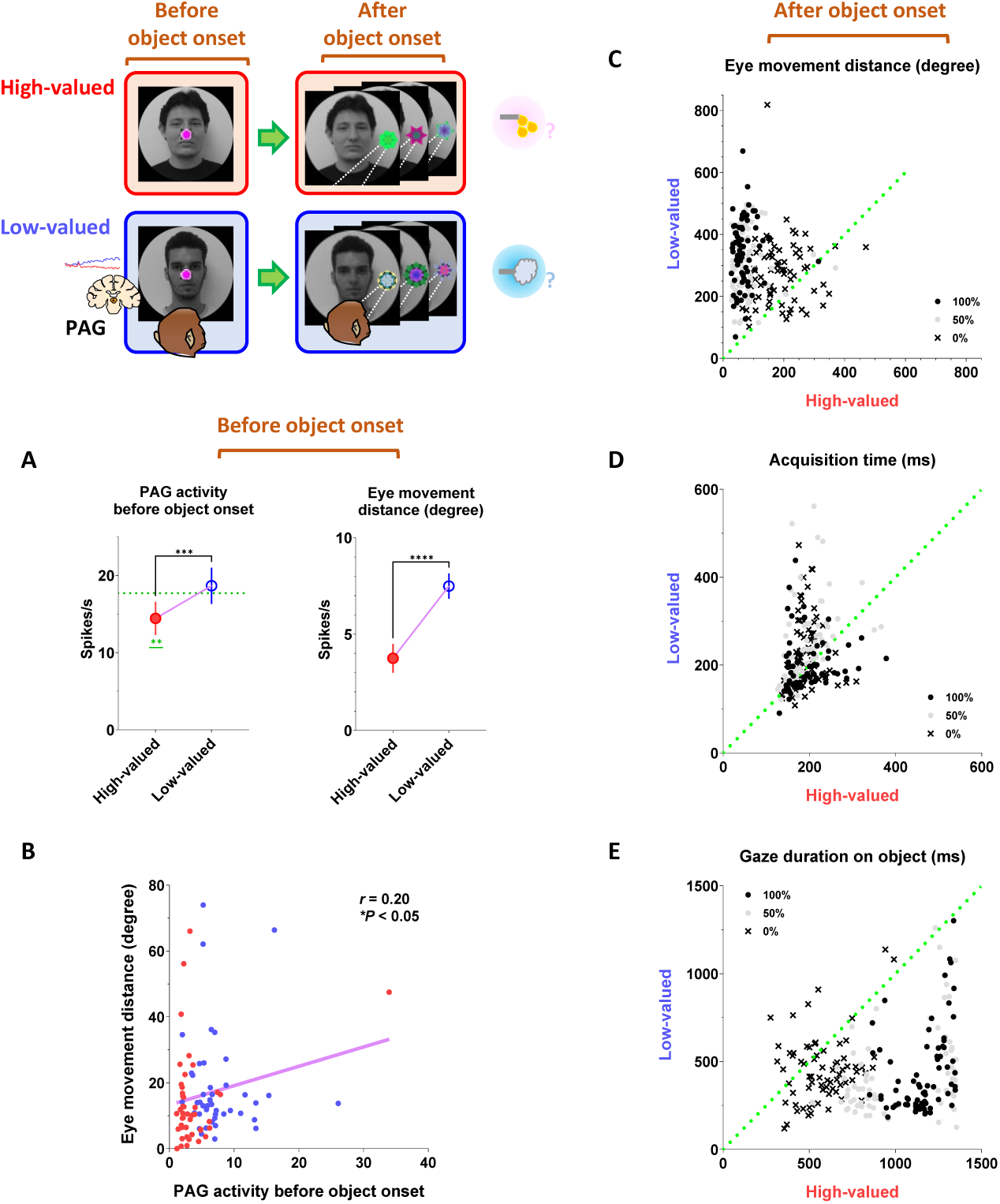
Implications of tonic PAG activity on subsequent behaviors through the inhibitory control of eye movements. (A) Left, the average firing rates of PAG neurons before object onset during the Pavlovian task. The firing rates were quantified for PAG neurons, as shown in figures 5O-P, during the - 200– 0 ms period preceding object onset. Right, the average eye movement distance before object onset in the Pavlovian task. The distances were quantified during the - 200–0 ms period preceding object onset (Wilcoxon matched-pairs signed rank test; *\*\*\*P* < 0.001, *\*\*\*\*P* < 0.0001, *n* = 44). (B) The correlation between the average firing rates of PAG neurons (from A, left) and the average eye movement distances (from B, right) (Pearson correlation analysis; *r* < 0.05, \**P* < 0.05, *n* = 44). (C) The average eye movement distance after object onset in the Pavlovian task. The distances were quantified for 1 s after object onset, as shown in figures A-D. (D) The time to initiate gaze on the object after object onset in the Pavlovian task. The start time for the monkeys’ gaze reaching the object was analyzed. (E) The duration of gaze fixation on the objects after object onset in the Pavlovian task. The gaze-holding duration was measured for 1.5 s.

Even after the object appeared, the differences in eye movements between the high-valued and low-valued scenes persisted and were sustained until the end of the trial (Fig. 6C). Thus, when the object subsequently appeared in the Pavlovian task, the monkeys in the low-valued scenes experienced slowness in locating their gaze on the object (Fig. 6D), and even after making a saccade, their gaze did not remain fixated as long as it did in the high-valued scenes (Fig. 6E).

## Discussion

### Tonic PAG activity and remaining reward expectancy in overcoming disappointment

This study proposes that when animals encounter disappointing events, they could overcome disappointment and continue reward-seeking behaviors through a balance between distinct phasic and tonic signals from the LHb and PAG.

The LHb is well-known as one of the primary inputs to the substantia nigra pars compacta (SNc) and ventral tegmental area (VTA) dopamine neurons for signaling disappointment.^14,43–46^ This neuronal pathway is essential for adaptability, flexibility, and learning of animal behaviors as it can modulate neuronal plasticity in the basal ganglia via dopamine transmission.^47–50^ The LHb responds to disappointing events with phasic firings based on the concept of reward prediction error, which calculates the discrepancy between actual outcomes and predicted values.^21,51–53^ Consequently, when an animal obtains a reward larger than expected, it suppresses LHb activity, enhances dopamine release to the striatum, and activates the direct pathway within the basal ganglia system. This pathway projects to the superior colliculus (SC), thalamus, and pedunculopontine nucleus (PPN), thereby facilitating the associated actions via the substantia nigra pars reticulata (SNr) and the internal segment of the globus pallidus (GPi).^54–56^ Conversely, when the reward is smaller than expected, LHb activity is stimulated, leading to reduced dopamine release and activation of the indirect pathway, which includes the external segment of the globus pallidus (GPe) and the subthalamic nucleus (STN), thereby suppressing the associated actions.^57–61^

However, to optimize reward acquisition, animals often need to overcome disappointment and sustain reward-seeking behaviors by considering additional factors (e.g., good or bad) beyond these prediction errors (i.g., better or worse). For instance, during reward-seeking behaviors, animals may encounter a disappointing event that is relatively worse than expected but could still hold an absolute value worth pursuing.^62^ Our findings revealed distinct phasic and tonic neuronal responses in the LHb and PAG, representing disappointment and remaining reward expectancy, respectively. Notably, the PAG activity was proficient at signaling reward expectancy throughout task trials, persisting until the desired reward was achieved (Fig. 4M-P and 5M-P). Meanwhile, despite occasional disappointments, the monkeys consistently engaged in a series of task procedures with heightened visual attention (Fig. 4A-D and 5A-D).

On the one hand, the PAG has reciprocal connections with the LHb, which should not be overlooked as a potential node for exchanging tonic signaling between them and guiding animals to overcome disappointing events.^63–67^ On the other hand, the PAG also establishes neuronal networks with other critical brain regions in the reward system, such as the ventral pallidum^68,69^ and amygdala (AMY).^70–73^ In previous studies, we particularly reported similar tonic activity patterns in amygdala neurons, which facilitated extraordinary goal-directed eye movements within the range of express saccades.^74,75^ Therefore, we suggest that this tonic signaling from the amygdala could be a strong candidate for modulating dopamine activity via the amygdalo-nigral pathway,^76,77^ and dopamine neurons could, in turn, integrate tonic signals from the PAG and phasic signals from the LHb (Fig. 7). Finally, further studies on the reciprocal connections of the PAG with the LHb and amygdala could deepen our understanding of how animals overcome disappointment and sustain reward-seeking behaviors, ultimately achieving desired outcomes. Indeed, the PAG has recently been highlighted as a potential therapeutic target for major depressive disorder,^78–80^ which could expand treatment approaches and complement LHb studies aimed at improving sustained antidepressant effects.^81–83^

**Figure 7.**
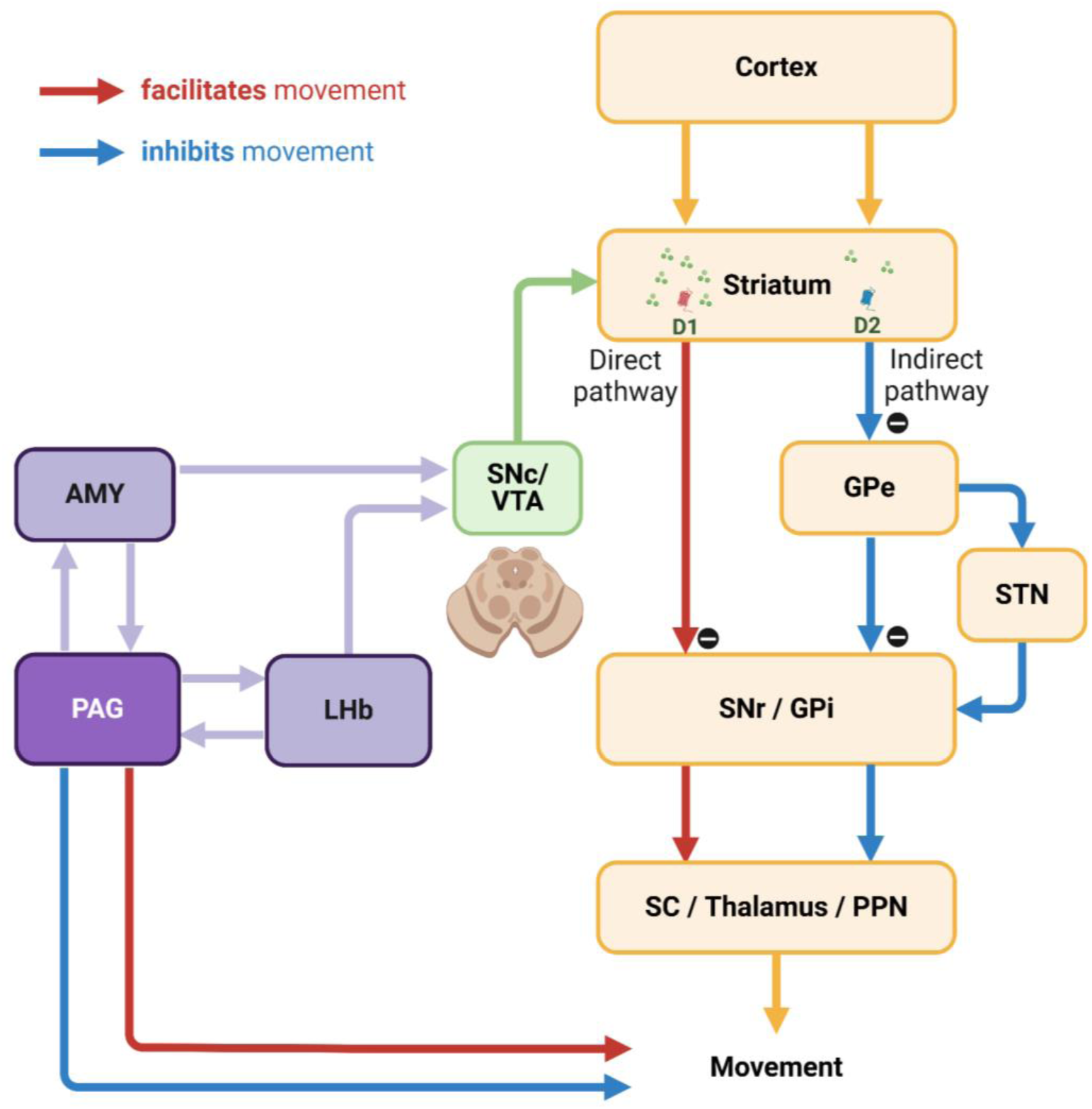
Hypothetical neuronal mechanisms underlying inhibitory control of movements. The interconnection between the PAG and AMY could play a key role in the inhibitory control of movements via dopamine neurons in the SNc and VTA. This pathway could enable dopamine neurons to integrate tonic signals from the PAG with phasic signals from the LHb. Furthermore, the PAG could be essential for integrating emotional information from the AMY and LHb, thereby orchestrating movement control through additional interactions with brain regions such as the forebrain, cerebellum, and other brainstem areas.

### Tonic suppression of PAG activity in bridging sequential reward-seeking behaviors

Next, our findings suggest that the tonic activity of the PAG could play a crucial role in bridging sequential motor actions involved in reinforcement learning. This is evident in the correlation between the tonic signaling of the PAG and the anticipatory gaze of monkeys, which consistently maintained their gaze on the background scene images where their tasks took place (Fig. 1E-F). Thereby, the anticipatory gaze enabled monkeys to prepare for subsequent actions (Fig. 3D-E and H-I), allowing them to detect the FP with greater speed and gaze rates (Fig. 3).

This is an essential component in the motor skill behaviors of animals, as it allows a sequence of actions to be interconnected and executed as cohesive behavioral units (i.g., motor chunk).^84–86^ As a result, animals could improve the speed and accuracy of a series of sequential procedures in reward-seeking behaviors.^87,88^ This is exemplified by our previous study, which examined motor skill acquisition in monkeys using a 2×5 task.^89^ The 2×5 task involved training the monkeys to learn and remember the sequence of pressing five pairs of buttons, which illuminated from a total of 16 buttons, in a predetermined order. Following a series of training sessions, the monkeys exhibited skilled behavior in which, upon pressing one button, they were able to locate their gaze on the next button they would press before it illuminated. As a result, they could perform subsequent actions with exceptional speed and accuracy, leading to faster reward acquisition.

Furthermore, this reinforcement between behavioral units could also play a pivotal role in gating subsequent behaviors sequentially. For example, monkeys initiate eye contact before engaging in social behaviors like lip-smacking.^90^ Sequentially, lip-smacking induces synchronization of the animal’s behavior, facilitating emotional exchange and cooperative actions, which, in turn, promotes a series of various subsequent behaviors.^91–93^ Thus, the tonic signaling of the PAG, which supported the monkeys’ anticipatory gaze toward face scene images (Fig. 1), could play a crucial role in enhancing the likelihood of animals engaging in subsequent social behaviors. These motor skills, including social behaviors, are important sources of natural rewards and pleasure in the lives of humans and animals.^94–96^ In this way, social interactions between animals have been shown to improve stress coping,^97–99^ mitigate aversive responses to unpleasant stimuli, and enhance the persistence of ongoing motivated behaviors.^100–104^ Therefore, from the perspective of reinforcement learning, we suggest that the tonic activity of the PAG can provide additional reward resources for overcoming disappointing events and continuing ongoing motivated behaviors by facilitating the connection between sequential motor actions.

### Tonic PAG activity in inhibitory control of unnecessary movements

Lastly, we suggest that tonic PAG activity can contribute to the inhibitory control of unnecessary movements, which could potentially implicate its significant role in movement disorders. In the present study, we observed that monkeys exhibited strong inhibition of unnecessary eye movements during highly motivated anticipatory scene viewing (Fig. 4E-H and 5E-G). The inhibitory motor control can play a crucial role in regulating impulsivity, guiding action selection, and prioritizing the sequence of actions in human and animal behavior.^105–107^ Thereby, this process ultimately improves the speed and accuracy of desired behaviors and smooths out the motion of humans and animals in response to the overwhelming influence of information processed through various sensory inputs that flow into the brain.^108–111^

These findings could further provide valuable insights into the implications of PAG activity in movement disorders, such as Parkinson’s disease. Parkinson’s disease is characterized by abnormalities in the initiation and speed of actions (e.g., akinesia and bradykinesia).^48^ On the other hand, another significant symptom observed in this disease is the impairment in suppressing unnecessary movements, such as tremors and dyskinesia. These symptoms manifest abnormalities not only in limb movements but also in eye movements.^112–114^ For example, a Parkinson’s disease patient in a previous study exhibited frequent abnormal repetitive saccades and fixation patterns during word reading, leading to more frequent unnecessary eye movements compared to a healthy individual.^115^ As a result, this impaired overall reading speed and comprehension.

A key characteristic of Parkinson’s disease is the degeneration of dopaminergic neurons in the SNc. Additionally, the PAG, along with surrounding brainstem regions, has also been reported as an area showing significant changes in patients and animal models with Parkinsonian symptoms.^32,37,116–120^ However, there is still a lack of research on the role of the PAG in Parkinsonian symptoms. The dopamine deficits in Parkinson’s patients have primarily been reported in the caudal dorsal lateral part of the SNc, which projects to the tail of the striatum (i.e., the caudal part of the striatum).^121–124^ This neuronal pathway from the caudal dorsal lateral part of SNc to the tail of the striatum plays a crucial role in regulating automatic movements.^125–129^ Consequently, neurodegeneration in this pathway can impair the function of the indirect pathway, which is responsible for the inhibitory control of unnecessary automatic movements,^130–132^ and the decrease in PAG activity may affect its interaction with these basal ganglia circuits.

Importantly, the PAG is also recognized as a critical integrator of emotional responses and outputs in the emotional motor system, projecting to the spinal cord, forebrain, cerebellum, and other brainstem regions to control movement (Fig. 7).^133–136^ For instance, dysfunction of the PAG might impair eye movement control in Parkinsonism through its connections, distinct from basal ganglia output, to the brainstem areas associated with the oculomotor system.^137–139^ Future studies investigating the specific mechanisms of PAG activity and its interplay with basal ganglia circuits could significantly enhance our understanding of the pathophysiology of movement disorders and identify new therapeutic targets.

## Materials and Methods

### General Procedures

Two male Macaca mulatta monkeys were used for this study. (CH, 10-year-old, 15 kg and KI, 10-year-old, 9.5 kg). All animal care and experimental procedures were approved by the Animal Care and Use Committee of the National Eye Institute and complied with the Public Health Service Policy on the Humane Care and Use of Laboratory Animals. Apple juice (high-value, 600 µl; low-value, 200 µl) and airpuff (10-20 psi, 100 ms) were used as reward and punishment outcomes in the task. Partial data from the LHb and behavioral recordings were used in related publications that tested the same task protocol.^21,92^ The monkeys’ eye positions (*n* = 72)were recorded using an EyeLink 1000 Plus eye tracker (SR Research) and were simultaneously recorded with neuronal signals via Blip software (www.robilis.com/blip/).

### Scene-based foraging/Pavlovian task

A trial of the scene-based foraging/Pavlovian task began with the appearance of a scene image (size: 40°; (2 faces + 2 landscapes) × 4 scene groups = 16 scene images per block), which was maintained as the background scene throughout the trial (Fig. 1B). After 1 s of free viewing, a FP indicating either the foraging or Pavlovian task appeared at the center of the scene, and the respective task began. Each block of the task consisted of 384 trials, with tasks presented in a pseudo-random order (foraging task, 192 trials; Pavlovian task, 192 trials). Additionally, 32 non-cued free outcomes (8 high-value rewards, 8 low-value rewards, 16 punishments) were randomly delivered between task trials without any visual stimuli.

### Background scene images

We created four groups of background scene images, each contextually associated with different reward and punishment outcome experiences (Supplementary Fig. S1): high-value reward without punishment (Rwd++ Pun-), high-value reward with punishment (Rwd++ Pun+), low-value reward without punishment (Rwd+ Pun-), and low-value reward with punishment (Rwd+ Pun+). This study focused on data from high-valued scenes (Rwd++ Pun-) and low-valued scenes (Rwd+ Pun+) to address the primary research question (Supplementary Fig. S1). As a result, monkeys received a large amount of juice reward for completing foraging tasks in the high-valued scenes and a small amount of juice reward in the low-valued scenes (Fig. 1A). During Pavlovian tasks, in the high-valued scenes, monkeys received rewards of the same size as those in the foraging tasks. In contrast, in the low-valued scenes, monkeys received airpuff punishment outcomes instead of juice rewards. The background scene images were collected from Google Earth (https://www.google.com/earth), OpenAerialMap (https://openaerialmap.org), and Face Database (https://fei.edu.br/~cet/facedatabase.html).

### Foraging task

The foraging task began with the appearance of a square-shaped FP (size, 2°), requiring the monkeys to fixate on it for more than 700 ms within 1 s (Fig. 4). After the fixation, one of the good or bad objects (size, 10°) appeared on either the left or right side (15°) of the background scene image. Monkeys were then required to fixate on the good object (500 ms) within 1 s to receive a reward. If the bad object appeared after fixation, monkeys had to avoid it by either not making a saccade to the object for 1 s or by fixating on the bad object for no more than 500 ms. If the monkey successfully avoided the bad object, it disappeared from the scene, and the FP reappeared at the center of the scene. The monkey was then required to fixate on the FP again. Afterward, the good object was presented again on either side of the scene, and the trial could be completed by fixating on the good object to acquire the reward. If the monkey failed to avoid the bad object or failed to fixate on the FP or the good object, the visual stimuli disappeared, accompanied by a beep sound. The trial was then repeated from the scene onset until the monkey correctly completed the task and acquired the reward. Therefore, monkeys could earn fixed amounts of reward outcomes for each scene image in the block. The good object appeared first in 1/3 of the trials and after the bad object in 2/3 of the trials. Each scene image had different fractal images^129^ for the respective good and bad objects.

### Pavlovian task

The Pavlovian task began with the appearance of a circular FP, which required no action from the monkeys for 1 s (Fig. 5). After 1 s, one of three objects (100%, 50%, or 0% probability) appeared on the left or right side of the background scene image for 1.5 s. Monkeys then received a reward or punishment outcome based on the probabilities associated with each object. One second after the outcome delivery began, the background scene disappeared, and the next trial started after approximately 7 s. Each scene image had different fractal objects representing the respective objects.

### Electrophysiology

We recorded 34 neurons from the LHb and 44 neurons from the PAG in two monkeys: CH (LHb, 15; PAG, 30) and KI (LHb, 19; PAG, 14). Single-unit neuronal activity was recorded using glass-coated electrodes (diameter 0.38 mm, 1 MΩ, Alpha-Omega) connected to a microelectrode AC amplifier (model 1800; A-M Systems; gain, 10k; filters, 0.1 to 10 kHz) and a band-pass filter (model 3384; Krohn-Hite). The electrode was advanced using an oil-driven micro-manipulator (MO-97A, Narishige) through a guide tube and an 8° tilted posterior chamber. Recording sites were confirmed via vertical MRI scanning (4.7 T, Bruker) with a gadolinium-filled grid (1 mm-spacing) and Elgioy deposits marking the PAG.^140^ Neuronal firing was monitored in real-time and isolated using custom voltage-and time-based windows in Blip software.

### Statistical Analysis

We presented data as mean ± standard error of the mean. The statistical significances were analyzed using the Wilcoxon matched-pairs signed rank test, one-way repeated measures analysis of variance (ANOVA) with Bonferroni post hoc test, and Pearson correlation analysis using Prism9 (GraphPad Software). The average firing rates of neurons and gaze probabilities were smoothed by Gaussian kernel (σ = 10 ms) using MATLAB (MathWorks).

## AUTHOR CONTRIBUTIONS

Conceptualization, H.L. and O.H.; Methodology, H.L. and O.H.; Software, H.L. and O.H.; Formal Analysis, H.L. and O.H.; Investigation, H.L. and O.H.; Writing –Original Draft, H.L.; Writing – Review & Editing, H.L. and O.H.; Visualization, H.L. and O.H.; Supervision, O.H.; Project Administration, H.L. and O.H.; Funding Acquisition, O.H.

## ACKNOWLEDGMENTS

We thank M.K. Smith, A.M. Nichols, T.W. Ruffner, D. Yochelson, A.V. Hays, J.W. McClurkin, J. Fuller-Deets, and current and former colleagues of LSR for discussions and technical support; D. Parker, H. Warnock, G. Tansey for animal facility assistance; S. Hong for providing recording software; S. Yamamoto, D.A. Leopold, K. W. Koyano, C.E. Thomaz, and his colleagues for providing visual stimuli; Kenji W. Koyano again for technical support for Elgiloy deposit marking, This research was supported by the Intramural Research Program at the National Institutes of Health, National Eye Institute (1ZIAEY000415).

## DECLARATION OF INTERESTS

The authors declare no competing interests.

## Notes

### Competing Interest Statement

The authors have declared no competing interest.

