## Supplementary Fig. S1 for "Periaqueductal gray passes over disappointment and signals continuity of remaining reward expectancy"

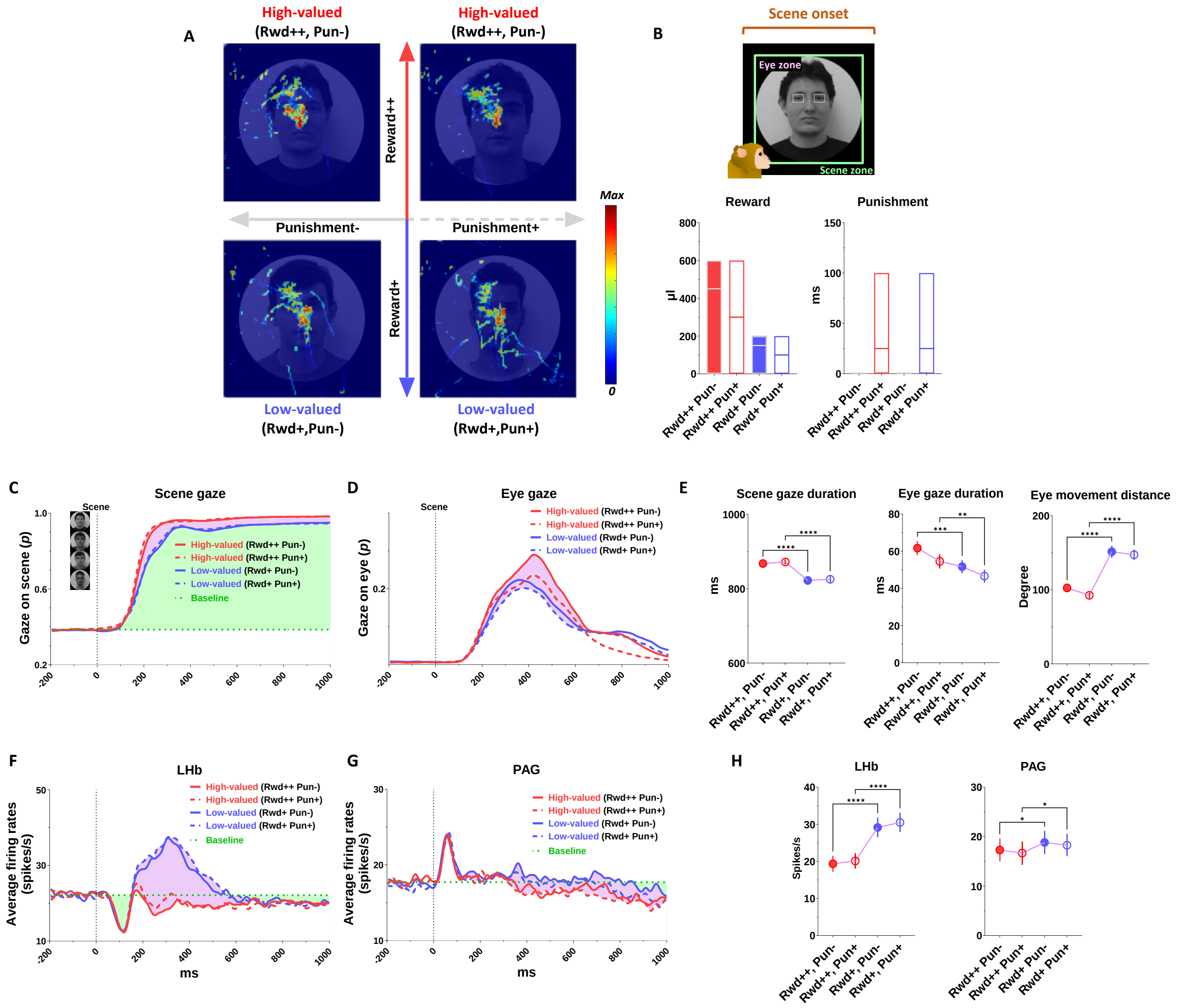


Supplementary Fig1

**Supplementary Fig. S1 Neuronal responses and anticipatory gazes were predominantly regulated by expectancy of reward rather than punishment**

(A) An example heatmap illustrating monkeys' gaze on face scene images. We conducted a scene-based foraging/Pavlovian task with four groups of background scene images: two high-valued scenes that provided a high-value reward, either without punishment (Rwd++ Pun-) or with punishment (Rwd++ Pun+), and two low-valued scenes that provided a low-value reward, either without punishment (Rwd+ Pun-) or with punishment (Rwd+ Pun+).

(B) The bars represent the average amount of juice reward and airpuff punishment provided for each group of scenes during the task procedures.

(C) The probabilities of gaze on face scene images during the free-viewing period. The purple shaded areas represent the differences in the responses between high-valued and low-valued scenes. The green shaded areas represent changes in the responses relative to the baseline during the task procedures.

(D) The probabilities of gaze on the eye regions of face scene images during the free-viewing period.

(E) Left, the duration of gaze on face scene images was quantified for 1 s during the free-viewing period. Middle, the duration of gaze on the eye zone was quantified for 450 ms after scene onset. Right, the distance of eye movements was quantified for 1 s during the free-viewing period (One-way repeated measures ANOVA with Bonferroni post hoc test; **$P < 0.01$, ***$P < 0.001$, ***$P < 0.0001$, $n = 72$).

(F, G) The average firing rates of LHb and PAG in response to scene onset.

(H) The average firing rates of LHb ($n = 34$) and PAG ($n = 44$) neurons in response to the scene onset. The LHb response was quantified during the 150–600 ms after object onset. The PAG response was quantified during the 250–800 ms after object onset (One-way repeated measures ANOVA with Bonferroni post hoc test; *$P < 0.05$, ***$P < 0.0001$).
